# Coevolution of the *Tlx* homeobox gene with medusa development (Cnidaria: Medusozoa)

**DOI:** 10.1101/2022.03.08.483504

**Authors:** Matthew Travert, Reed Boohar, Steven M. Sanders, Matthew L. Nicotra, Lucas Leclère, Robert E. Steele, Paulyn Cartwright

**Author notes:** Department of Biology, University of Miami, Coral Gables, Florida (Reed Boohar). Matthew Travert, **Email:**. **Author Contributions:** Paulyn Cartwright, Steven Sanders, Robert Steele, Matthew Travert designed research; Reed Boohar, Steven M. Sanders and Matthew Travert performed research; Lucas Leclère, Matthew L. Nicotra, Steven M. Sanders and Matthew Travert contributed new reagents/analytic tools; Paulyn Cartwright, Lucas Leclère and Matthew Travert analyzed data; Paulyn Cartwright and Matthew Travert wrote the paper with input from the authors.

## Abstract

The jellyfish, or medusa, is a life cycle stage characteristic of the cnidarian subphylum Medusozoa. By contrast, the other cnidarian subphyla Anthozoa and Endocnidozoa lack a medusa stage. Of the medusozoan classes, Hydrozoa is the most diverse in terms of species number and life cycle variation. A notable pattern in hydrozoan evolution is that the medusa stage has been lost or reduced several times independently. Although this loss of the jellyfish stage is thought to be due to heterochrony, the precise developmental mechanisms underlying this complex pattern of medusa evolution are unknown. We found that the presence of the homeobox gene *Tlx* in cnidarian genomes is correlated with those medusozoans that have a medusa stage as part of their life cycle. Although *Tlx* is conserved in Bilateria and Cnidaria, it is missing in the genomes of anthozoans, endocnidozoans, and those hydrozoans that have lost the medusa stage. Selection analyses of *Tlx* across medusozoans revealed that hydrozoans undergo relatively relaxed selection compared to the other medusozoan classes, which may in part explain the pattern of multiple medusa losses. Differential expression analyses on three distantly related medusozoan representatives indicate an upregulation of *Tlx* during medusa development. In addition, *Tlx e*xpression is spatially restricted to regions of active development in medusae of the hydrozoan *Podocoryna carnea*. Our results suggest that *Tlx* plays a key role in medusa development and that the loss of this gene is likely linked to the repeated loss of the medusa life cycle stage.

## Introduction

Jellyfish represent part of a complex life cycle that is characteristic of the cnidarian subphylum Medusozoa, which includes true jellyfish (Scyphozoa), box jellyfish (Cubozoa), stalked jellyfish (Staurozoa) and hydromedusae (Hydrozoa). Medusozoans possess a metagenetic life cycle alternating between an asexual phase in the form of a sessile polyp and a sexually reproducing, typically pelagic, phase called the medusa or jellyfish. The medusa exhibits several distinct features including a bell-shaped morphology, striated muscles, gonads, and sensory organs. The parasitic Endocnidozoa is sister to Medusozoa (Chang et al., 2015) and lacks a definitive polyp or medusa stage. The other major cnidarian class, Anthozoa (corals, anemones and sea pens) also lacks a medusa stage as well as all traits associated with this free-living stage, and instead possesses a monogenetic life cycle, comprising only sexually and asexually reproducing sessile polyps. Given that these other two major classes lack medusae characters, it can be assumed that medusae are a derived feature of Medusozoa and not ancestral to Cnidaria.

Despite the medusa being characteristic of Medusozoa, losses of this life cycle stage within Hydrozoa are omnipresent and most likely are due to developmental heterochrony (Boero and Bouillon 1987; Boero et al., 1992; Kubota, 2000). The fully developed hydromedusa exhibits distinctive features such as a muscular structure at the bell margin called a velum, striated muscles, marginal tentacles and tentacle bulbs, a manubrium which contains the gut and mouth, and a gastrovascular system composed of radial and circular canals. The truncation of the medusa stage correlates with the loss or reduction of the aforementioned features, with the varying degrees of medusa truncation often mirroring stages of medusa development (Hincks 1868, Allman 1872, Goette 1916 and Namikawa 1991). The differing degrees of medusa truncation across species is a type of paedomorphic progenesis (Gould 1985, Cunningham and Buss 1993), where somatic development is truncated due to earlier sexual maturation.

While the development of the hydrozoan polyp has been well studied, particularly in the model systems *Hydra* (Galliot, 2012) and *Hydractinia* (Frank et al., 2020), the molecular mechanisms underlying the development of the hydromedusa remain poorly understood. While some studies have indicated that medusa development co-opts key developmental pathways functioning in the polyp (Sanders & Cartwright, 2015a, b; Kraus et al., 2015, Masuda-Nakagawa et al., 2000, Müller et al., 1999), recent studies suggest that some medusa-specific transcription factors might act as molecular switches that regulate aspects of medusa development (Leclère et al, 2019 and Kahlturin et al, 2019). Amongst identified “medusa specific” genes a large proportion are homeobox genes, suggesting that these genes may play a key role in the development and maintenance of the medusa.

One purported “medusa specific gene”, is the T Cell Leukemia Homeobox gene (*Tlx*) (Leclère et al, 2019). In vertebrates *Tlx* (also referred to as *HOX11*) is involved in spleen organogenesis, brain and skeleton patterning and in *Drosophila melanogaster* it is involved in distal patterning of the leg *(clawless)* (Lenti et al., 2016, Kojima et al., 2005)*. Tlx* could not be found in the genome of the sea anemone *Nematostella vectensis* (Kamm and Schierwater, 2006) and thus was thought to be absent in cnidarians. Here we show that *Tlx* is indeed present in cnidarians. However, our survey of this gene across cnidarian genomes found that it is present only in those cnidarian lineages exhibiting a medusa stage. *Tlx* is absent in all anthozoans and endocnidozoan genomes surveyed. In addition, an intact *Tlx* is absent in nearly all hydrozoans surveyed that have lost a medusa life cycle stage. In several distantly related medusae-bearing species, we found that *Tlx* expression is upregulated during medusa development and its expression is consistent with it playing a role in medusae patterning.

## Results

### The cnidarian *Tlx* ortholog shares a highly conserved genomic structure with bilaterian *Tlx*

Cnidarian and bilaterian *Tlx* genes are remarkably conserved in structure, including an EH1 domain near the N-terminus, the homeodomain, and its signature motif, RRIGHPY just upstream of the homeodomain, called the N-terminal arm (Figure 1A). In our search of public databases for *Tlx*, the EH1 domain, as well as the N-terminal arm were invariably found in complete sequences of *Tlx* in both cnidarians and bilaterians. However, these conserved regions could not be found in the putative *Tlx* orthologs of sponges and placozoans, and although ctenophore sequences had an EH1 domain, they lacked the N-terminal arm. In addition to low sequence identity in the homeodomain, a Bayesian phylogenetic analysis did not recover the putative *Tlx-like* gene previously identified in ctenophores (Pang and Martindale, 2008 and Derelle and Manuel, 2007), within the strongly supported bilaterian and cnidarian *Tlx* orthology group (Figure S1). This suggests that *Tlx* arose in the last common ancestor of Cnidaria + Bilateria. Vertebrates possess three *Tlx* paralogs, suggesting two duplication events in the last common ancestor of vertebrates. A phylogenetic tree consisting of 38 cnidarian taxa and 11 vertebrate taxa including their three paralogs is shown in Figure 1B. *Tlx* forms a well-supported orthology group (BS=83, PP=0.99). Although vertebrate *Tlx* paralogy groups are respectively well supported, the relationship between cnidarian *Tlx* to a specific vertebrate paralog could not be recovered (TLX1/3) with sufficient support. A phylogenetic analysis including select protostome, cnidarian and vertebrate *Tlx* genes also formed a well support *Tlx* orthology group in the Bayesian analysis (PP=0.98) but failed to recover specific orthology relationships between major taxa (Figure S1).

**Figure 1.**
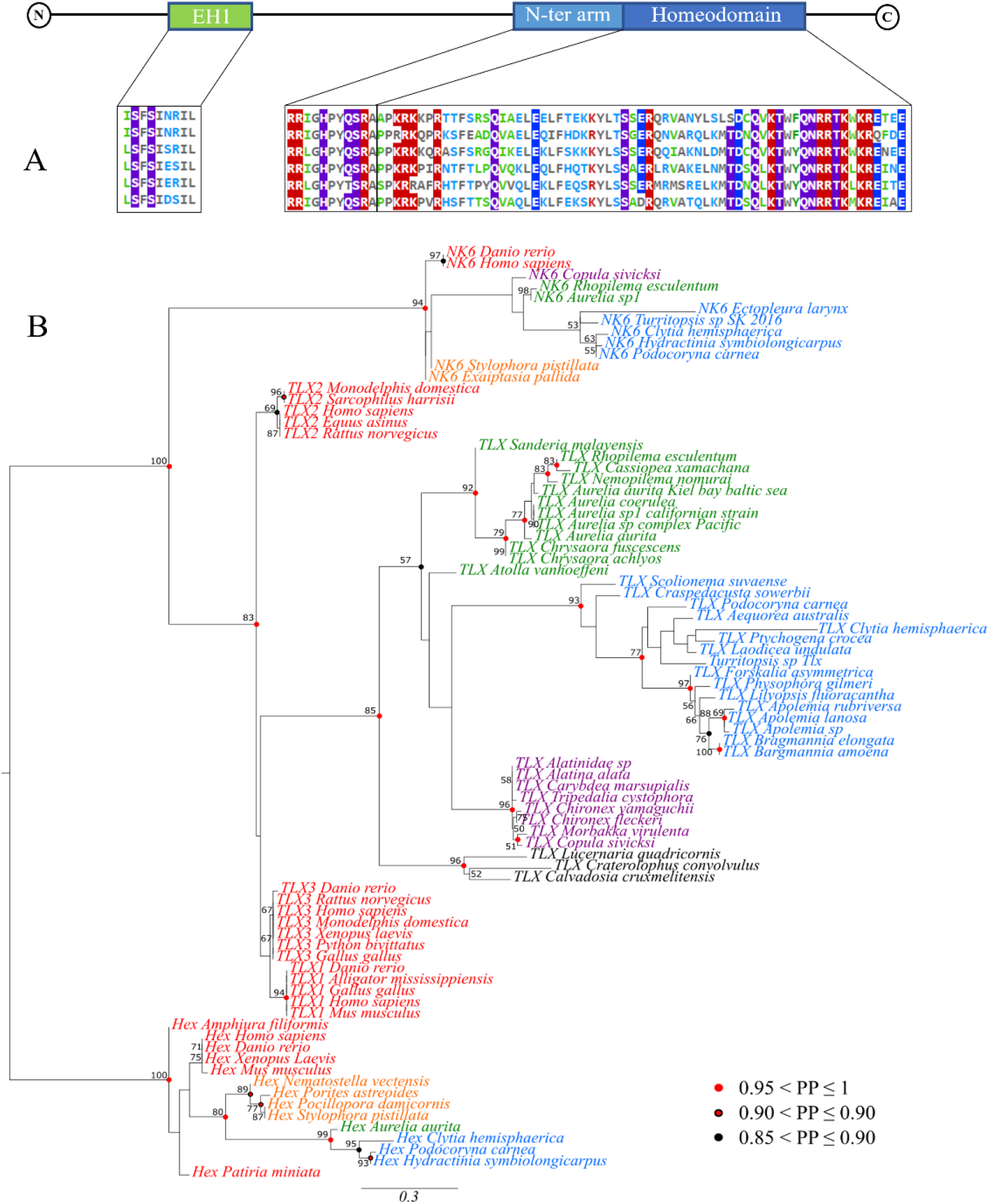
A) Schematic of TLX genomic structure and the corresponding amino acids sequence alignment for TLX conserved domains from six representative medusozoan taxa. Highly conserved positions are highlighted (>70% identity). Colors represent features of the position, purple (polar uncharged), red (positively charged), blue (negatively charged) and green (hydrophobic) B) Phylogram from maximum likelihood analysis, with three NK-L representatives (*Tlx*, *Nk6* and *Hex*). Vertebrate sequences are in red, hydrozoan sequences in blue, staurozoan sequences in black, scyphozoan sequences in green, cubozoan sequences in purple and anthozoan sequences in orange. Bootstrap values greater than 50% are indicated (1000 bootstraps) next to the nodes. Bayesian posterior probability greater than or equal to 85% are reported on the nodes with colored circles (color code on the figure). *Hex* sequences are used as the outgroup. Scale bar = number of inferred substitutions per position in the alignment.

### *Tlx* is absent from available genomes of cnidarians lacking a medusa

We define the presence of a medusa by the presence of medusa specific characters. Reduced medusae are often referred to as eumedusoids, cryptomedusoids or sporosacs (Bouillon et al., 2006) depending on their degree of developmental arrest. Here we call reduced medusae eumedusoids if they exhibit a gastrovascular system, velum and tentacles but lack discrete gonads and a mouth, cryptomedusoids if they bear only radial canals and highly reduced tentacle processes and sporosacs if they represent a fixed gonophore lacking any medusa features. Some hydrozoans, such as *Hydra*, do not bear any gonophores and instead release their gametes directly from the body column of the polyp. Given that eumedusoids possess many medusa-specific characteristics, we consider those species bearing eumedusoids as having a medusa stage, whereas those bearing cryptomedusoids, sporosacs or absence of any gonophore, we consider lacking a medusa stage.

In our search for the *Tlx* gene in 70 publicly available cnidarian draft genome assemblies we found that the presence of *Tlx* is invariably correlated with the presence of a medusa in the cnidarian life cycle. Specifically, *Tlx* was found in all 27 of the draft genomes from species that have medusae and not found in any of the 43 available draft genomes from species that lack medusae, including all anthozoans and endocnidozoans and the six hydrozoans that lost the medusae stage (Table 1). Although Ryan et al (2006) reported a *Tlx* gene in the sea anemone *Nematostella*, this gene lacks the signature EH1 domain and N-terminal arm and was not recovered in the *Tlx* orthology group with sufficient support in their analysis.

**Table 1.**
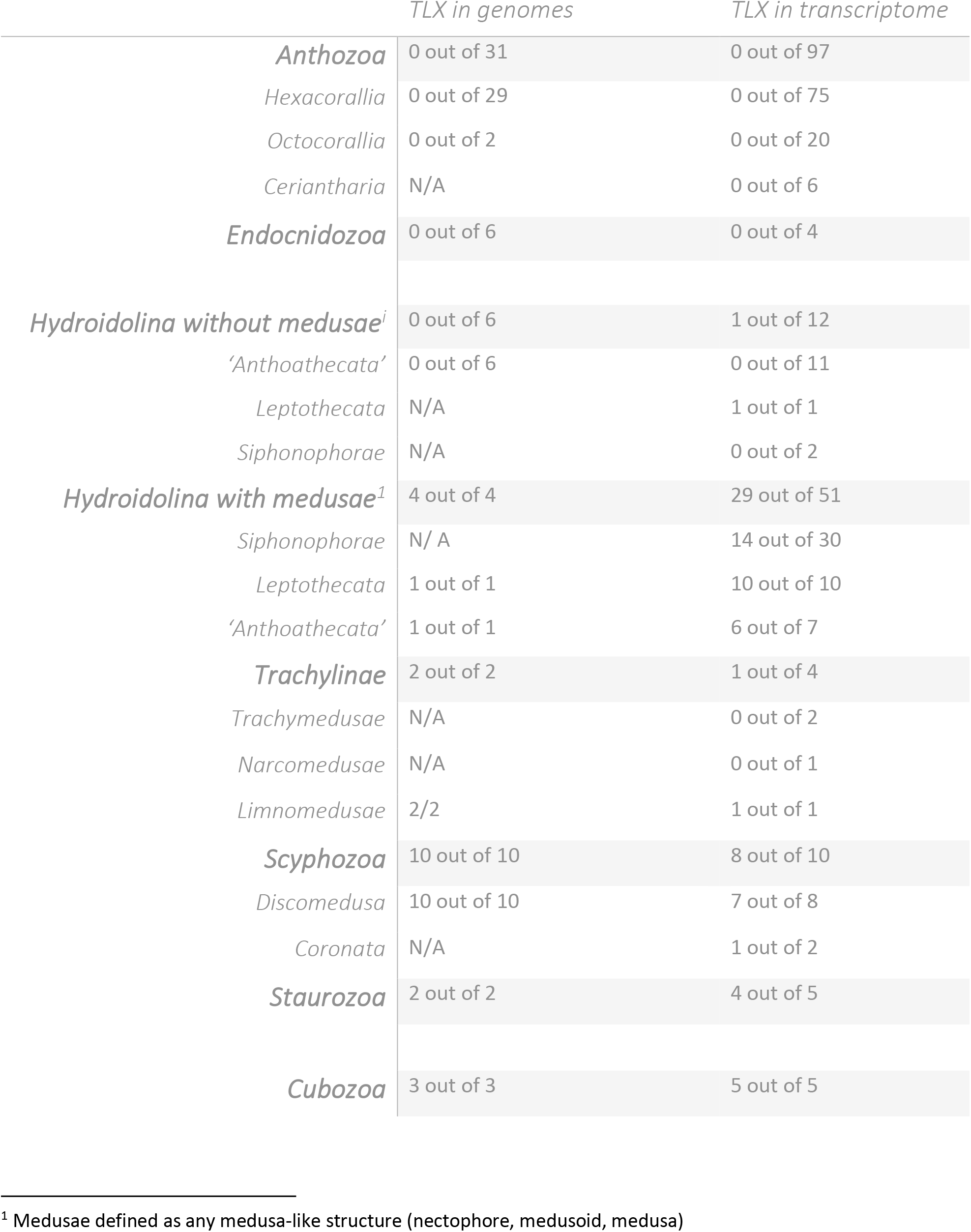
*Tlx* presence in publicly available cnidarian genomes and transcriptomes

*Tlx* was also searched for in 188 cnidarian transcriptomes, wherein the gene was present in 47 of 75 transcriptomes from medusa-bearing species (Table. 1). The failure to recover *Tlx* in some of the transcriptomes of medusa-bearing species does not necessarily indicate its absence from the genome, as transcriptomes are a limited subset of expressed genes of the tissue sampled. Therefore, the absence of *Tlx* in some of the sampled medusa-bearing species is likely the result of the particular tissue and/or developmental stage from which the transcriptome was generated. *Tlx* was not present in any available transcriptomes from species that lack a medusa, with three exceptions (*Millepora squarrosa, Ectopleura larynx* and *Dynamena pumila*) (Table 1). *Millepora squarrosa Tlx* is likely a pseudogene as it exhibits premature stop codons and the *Dynamena pumila* sequence was a partial sequence lacking the EH1 domain and the C-terminus of the homeodomain and thus the functionality cannot be inferred. *Ectopleura larynx* has the characteristic *Tlx* domains. It does however exhibit a unique codon insertion, leading to a threonine in the highly positively charged N-terminus of the homeodomain. Whether this change could affect *E. larynx Tlx* function is unknown.

Due to the sampling restrictions imposed by screening *Tlx* from publicly available cnidarian genomes, and the limitations of transcriptomes for estimating the presence of a gene as discussed above, we used degenerate PCR to screen genomic DNA from 100 medusozoan taxa for the presence of *Tlx*, including all medusozoan suborders, in order to span the breadth of medusozoan diversity. The primers were designed to amplify the N-terminal arm and the entire homeobox region. In the 69 taxa surveyed that have a medusa (or eumedusoid), a *Tlx* gene fragment was successfully amplified in 58 (84%). The failure to amplify a *Tlx* fragment in the other 11 medusa-bearing taxa could be due to the limitations of degenerate PCR, which is highly sensitive to DNA quality and primer binding. An amplification product was not obtained in 28 out of 31 taxa (90%) that lack a medusa (sporosac, cryptomedusoid or no gonophore). The three non-medusa bearing species for which a *Tlx* fragment was recovered were the cryptomedusoid bearing *Ectopleura larynx*, also found in the transcriptome above, as well as two species that bear sporosacs (*Amphisbetia minima* and *Sertularia perpusilla*). The sequence from *Sertularia perpusilla*, like the sequence of *Millepora squarrosa* discussed above, is likely a pseudogene as it contains several premature stop codons. Thus, of the total of five TLX sequences isolated from non-medusae bearing species from transcriptomes and/or PCR, only *Amphisbetia minima* has a typical TLX sequence.

### The absence of the *Tlx* gene is correlated with the absence of the medusa in Hydrozoa

Using the medusozoan phylogenetic tree from Cartwright and Nawrocki (2010) we pruned the taxa to match those samples for which degenerate PCR data were generated and reconstructed the evolution of the medusa stage. Similar to findings of Cartwright and Nawrocki (2010) our analysis inferred several independent instances of reduction and loss of the medusa stage. Out of the 14 such cases, seven are reductions to sporosacs, four to eumedusoid, one to cryptomedusoid and two are complete losses of the gonophore (Figure 2).

**Figure 2.**
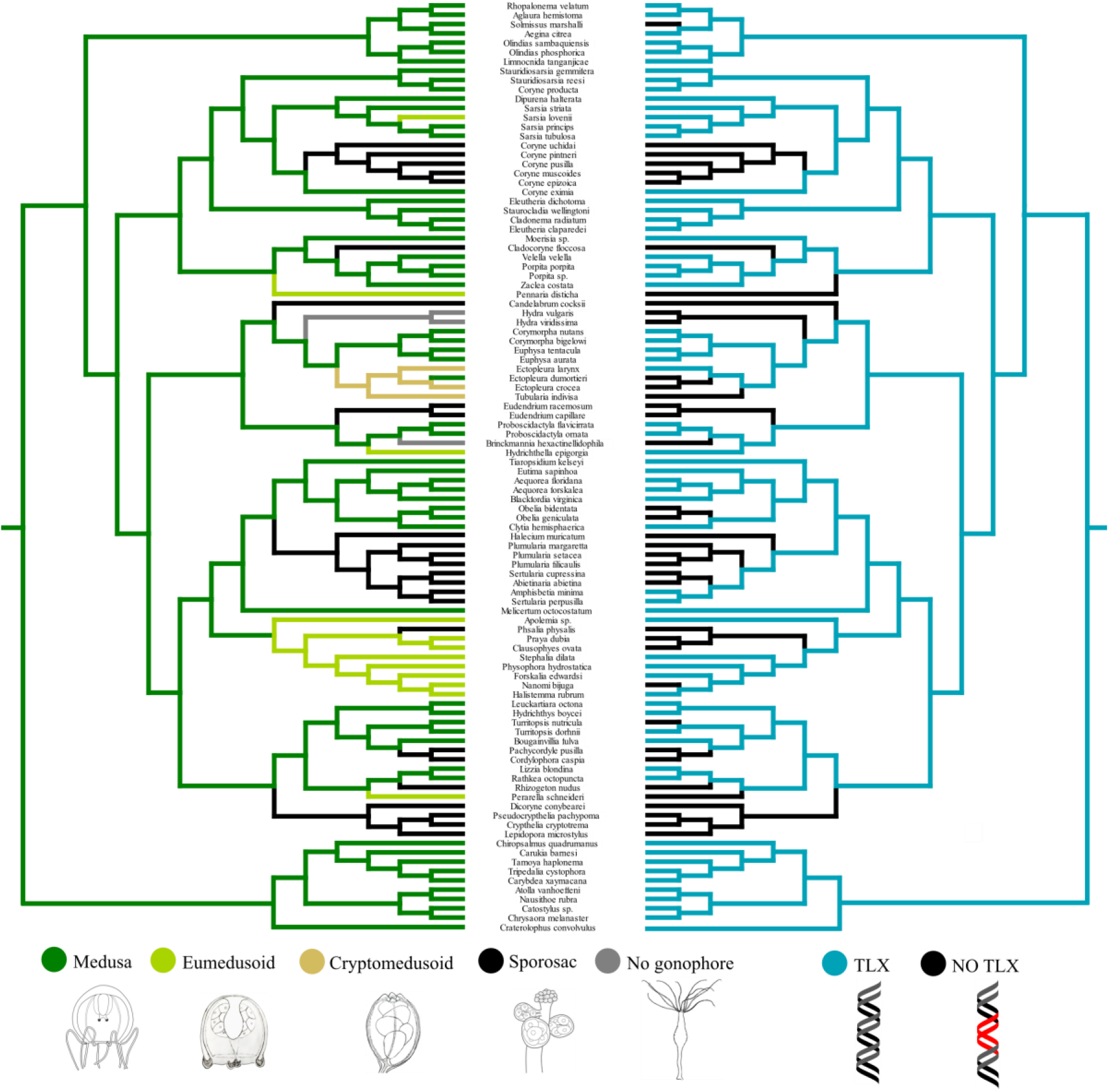
Wagner maximum parsimony ancestral character state reconstructions of medusozoan reproductive systems (left) against the Dollo maximum parsimony ancestral character state reconstructions of the presence of *Tlx* (right). Characters are unweighted and unordered, no optimization model was applied. Branches are colored in terms of degree of medusa reduction and presence of *Tlx* (shown in legend). Phylogeny pruned from Cartwright and Nawrocki, 2010 and rooted with Acraspeda.

A Bayesian correlation analysis of the presence of *Tlx* and the presence of medusa-like structures (medusa and eumedusoid) shows a very strong correlation between the two traits (Log Bayes factor = 19.535592). The same analysis also supports the presence of TLX together with the medusa stage as ancestral in Medusozoa (PP=1) (Figure 2).

### TLX homeodomain shows evidence of relaxed selection in species lacking medusa-like structures

To further investigate the apparent conservation of the TLX homeodomain amongst medusa bearing and the few non-medusa bearing lineages, we tested for relaxation/intensification of selection on the *Tlx* homeodomain in a codon-based phylogenetic framework. Using selection analyses, we tested the four *Tlx* sequences that were found from non-medusa-bearing species (*Millepora squarrosa*, *Ectopleura larynx*, *Dynamena pumila, Amphisbetia minima*) for relaxed selection. *S. perpusilla* sequence was removed from the analysis as the aberrant sequence did not allow for proper codon alignment. When testing these four sequences against the 46 medusa bearing reference species sequences, we found strong evidence for a relaxation of selection on *Tlx* in those medusa-less lineages (K=0.15. p=0.0000). While testing those four lineages independently, the same evidence of relaxation of selection was found except for *E. larynx* (K=1.19, p=0.1715) for which a non-significant intensification was inferred. By contrast, a significant intensification of selection on the *Tlx* homeodomain was detected for Acraspeda (Scyphozoa, Cubozoa and Staurozoa) (K=1.66, p=0.0000) and the hydrozoan order Siphonophorae (K=2.30, p=0.0050), using medusa-bearing hydrozoan species as a reference. No significant trend in selection was detected for the other hydrozoan lineages (see Table 2). This relaxation of selection in Hydrozoa may in part explain the pattern of multiple medusa losses that is not found in the other medusozoan classes.

**Table 2.** Analyses of the intensity of the selection on the TLX homeodomain within medusozoans

| Reference branches<br>Test branches | Medusozoans with medusae |  | Hydrozoans with medusae |
| --- | --- | --- | --- |
| Hydrozoans without medusae | K=0.15, p<0.0001 | Acraspeda | K=1.66, p<0.0001 |
| *Amphisbetia minima* | K=0.00, p=0.0010 | Trachylina | K=0.78, p=0.413 |
| *Dynamena pumila* | K=0.16, p<0.0001 | Siphonophorae | K=2.30, p=0.0050 |
| *Ectopleura larynx* | K=1.19, p=0.7710 | Leptothecata | K=1.26, p=0.521 |
| *Millepora squarrosa* | K=0.66, p=0.0249 | Anthoathecata | K=0.72, p=0.195 |
| Amplicons from degenerate PCR | K=0.71, p=0.1715 | Amplicons from degenerate PCR | K=1.04, p=0.917 |

Lastly a FUBAR test (Fast, Unconstrained Bayesian Approximation, Murrel et al., 2013) was performed to identify site specific variation in the selection of cnidarian TLX. Unsurprisingly, 9 codons in the EH1 domain and 75 codons in the homeodomain (including the N-terminal arm and flanking regions), respectively, showed evidence for pervasive purifying selection (PP>0.99), and no phylum-wide diversifying selection was detected upon remaining sites. Interestingly, the analysis detected smaller motifs flanking the homeodomain and its N-terminal arm. Amongst scyphozoans, a significant (PP>0.99) episodic purifying selection was detected on sites flanking the homeodomain (PWQILXK upstream of the N-terminal arm and TEEEKEEQRHAL downstream of the homeodomain), while a significant (PP>0.99) episodic purifying selection was detected in the flanking regions of the homeodomain of hydrozoan TLX, corresponding to a highly conserved CXC motif upstream of the N-terminal arm and a EINEMXEQQXR motif downstream of the homeodomain. Flanking motifs found in hydrozoans and scyphozoans do not provide direct information regarding the binding target of the homeodomain but could suggest that the target or the affinity for the target of TLX might differ between these lineages.

### Expression of *Tlx* is up-regulated during medusa development

To investigate the expression profile of *Tlx* in the medusozoan life cycle, we performed a differential expression (DE) analysis on distinct life cycle stages for the scyphozoan *Aurelia coerulea*, and two medusae-bearing hydrozoan species *Podocoryna carnea* and *Clytia hemisphaerica*. For *Aurelia*, *Tlx* expression is first detected in the polyp (scyphostoma), peaks when the polyp is producing medusae (strobilating) and is then downregulated in the juvenile medusa (ephyra) and mature medusa stages to expression levels comparable to the scyphostoma (Figure 3a). In *Clytia hemisphaerica, Tlx* expression is first detected at low levels during the planula stage and is maintained at low level in the feeding polyp (gastrozooid). *Tlx* expression is upregulated in the reproductive polyp that buds medusae (gonozooid) and the newly released juvenile medusa. In the adult medusa of *Clytia*, the expression of *Tlx* is downregulated to an intermediate level (Figure 3b). In *Podocoryna carnea* no significant expression of *PcTlx* is detected at the planula stage, the non-reproductive polyp nor in the reproductive polyp when the medusae buds are initially detected.

**Figure 3.**
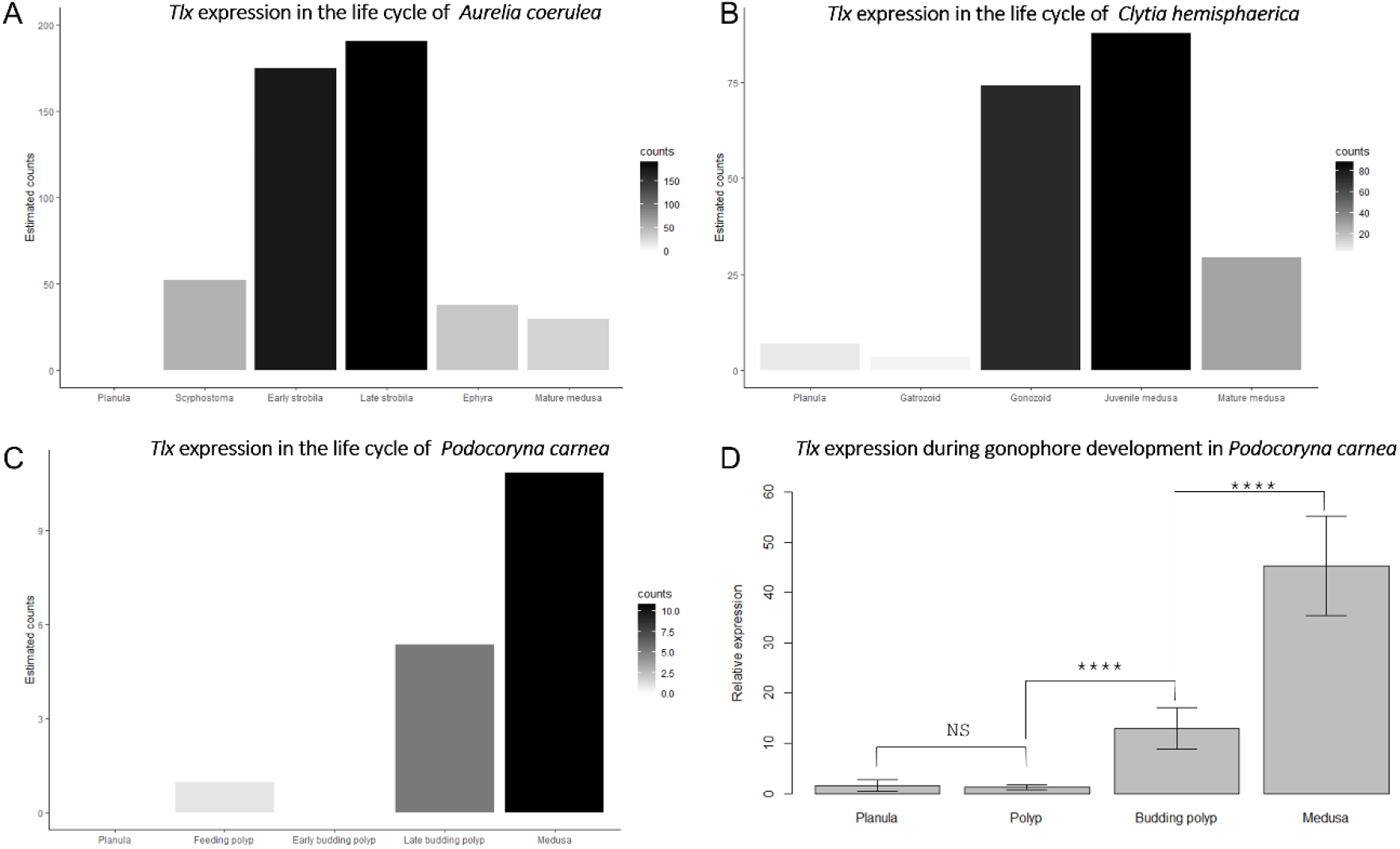
Normalized expression of *Tlx* in corrected counts for life cycle developmental stages (A-C). A) *Aurelia coerulea*,, B) *Clytia hemisphaerica*, C) *Podocoryna carnea*. The gradient charts indicate the breadth of *Tlx* corrected counts values for each species. D) RT-qPCR in *Podocoryna carnea*. (****) indicates a two tailed p-value <0.0001 from t-tests, (NS) indicates a non-significant two tailed p-value (p-value = 0.4638) from unpaired t-test.

*PcTlx* expression is upregulated in reproductive polyps budding medusae, during later stages of medusae development and remains at this expression level in the fully developed medusa after it is released from the polyp (Figure 3c). Although DE patterns were significant for the three species using estimated counts, other metrics, namely Transcripts per million (TPM) and Fragments Per Kilobase Million (FPKM), were inconclusive, likely due to the overall low expression of *Tlx*. To validate the RNA-Seq results in *Podocoryna*, we performed RT-qPCR on planulae, non-reproductive polyps, budding polyps and released medusae of *P. carnea* and found a significant difference in the relative expression of *Tlx* between the four life cycle stages (ANOVA, p<0.0001), as well as a higher expression of *Tlx* in released medusae compared to budding polyps (t-test, p<0.0001), (see Figure 3d).

### *Tlx* expression is spatially restricted during medusa development

*Tlx* expression was detected by whole mount *in situ* hybridization in juvenile and mature medusae of *Podocoryna carnea* but not in feeding and reproductive polyps (Figure 4A-C), or planulae (not shown). In *Podocoryna*, the gonads develop on the manubrium (the structure that contains the mouth and gut). *Tlx* expression was detected in an endodermal cell subpopulation surrounding the gonads of newly released medusae (Figure 4C, D) and additionally in the oocytes of female medusae (Figure 4F). This expression is initially localized at the base of the manubrium (not shown) in one day old medusae and expands mid-orally in the endoderm surrounding the gonads (Figure 4D). *Tlx* was sporadically detected in isolated ectodermal cells in the manubrium as well. In older medusae, the expression was also detected in the endoderm of the tentacle bulbs (structure on the bell margin proximal to the tentacles) and at the position of newly developing tentacles (Figure 4E).

**Figure 4.**
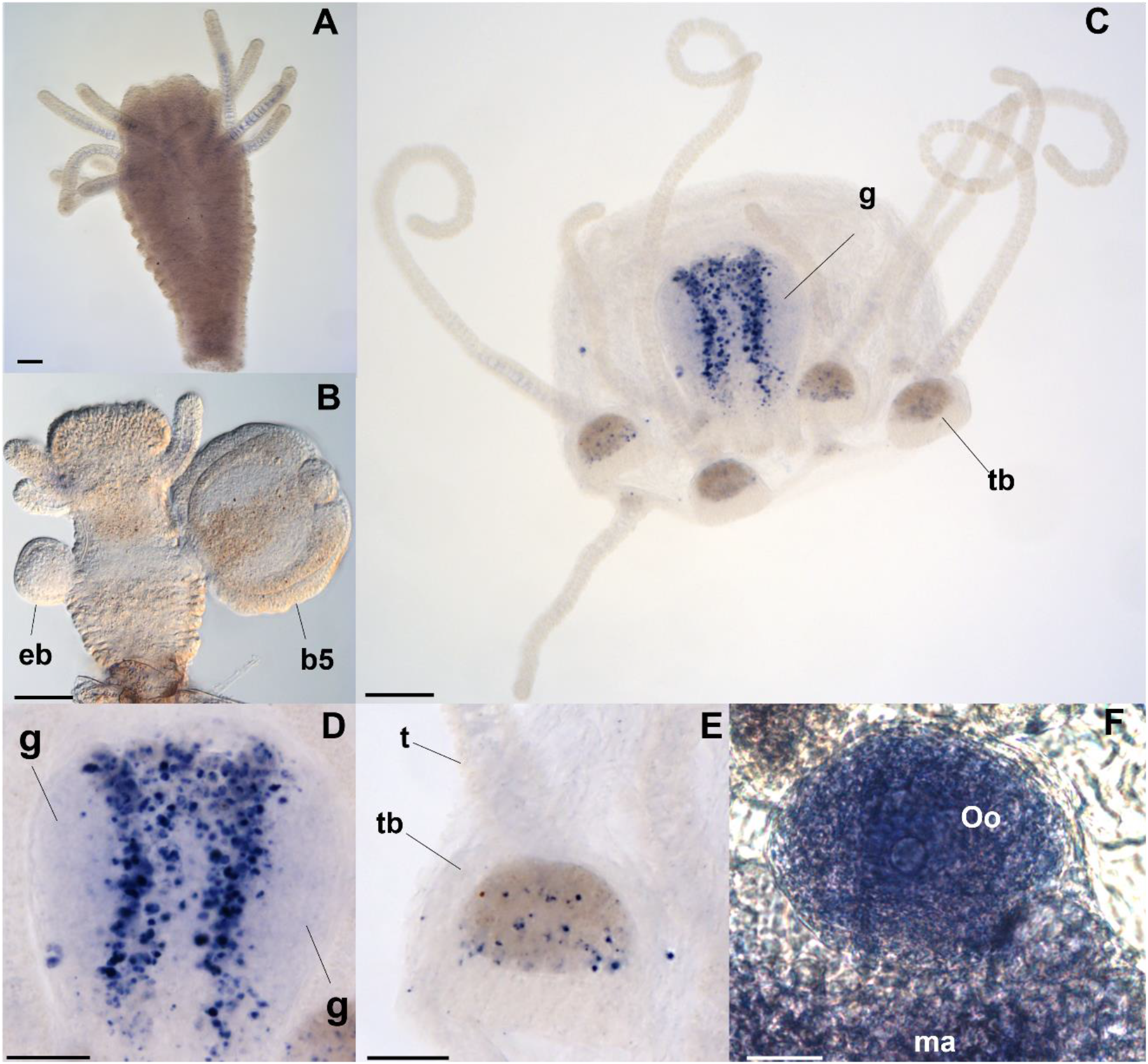
*In situ* localization of *PcTlx* transcript*s* on *Podocoryna carnea*. A) in male non-reproductive polyp, B) in a male budding polyp, C) in a male medusa, D) higher magnification of (A) of the manubrium, E) higher magnification of (A) of a tentacle bulb, F) and a mature oocyte in a female medusa, showing *Tlx* expression,. Samples are presented in radial view except the tentacle bulb that is presented in oral view. The manubrium (D) is presented oral down, aboral up. The tentacle bulb (G) is presented proximal left, distal right. Abbreviations: b5, medusa bud stage 5; eb, early bud (stage 1); g, gonad; t, tentacle; tb, tentacle bulb. Scale bar: 200 μm (A-C), 100 μm (D and E), 20μm (F).

## Discussion

Our detailed analysis of phylogenetic distributions of the medusa stage and the *Tlx* sequence reveals a striking correlation between the presence of an intact *Tlx* and the presence of the medusa life cycle stage in cnidarians. The few occurrences of *Tlx* in non-medusa bearing species are characterized by sequence alteration and/or relaxed selection in conserved regions, suggesting conversion to pseudogenes and possible loss of TLX function in those species.

In addition, our RNA-seq analyses show striking upregulation of *Tlx* during medusa development in three disparate medusozoans, suggesting the existence of a conserved role of *Tlx* in the development of the medusa, despite having very distinct developmental trajectories. Specifically, while the hydrozoan medusae relies on lateral budding from the polyp, *Clytia* possesses dedicated polyps (gonozooids) for budding medusae, whereas *Podocoryna* transforms its feeding polyp to a reproductive polyp upon onset of medusa budding. *Aurelia* does not bud medusae but undergoes a process of transverse fission of its polyp, called polydisc strobilation.

*PcTlx* expression is localized in cell populations distinct from the germline in areas of active somatic development, as well as mature oocytes in the released medusa. This suggests that *Tlx* plays a role in the maintenance of somatic development in the developing medusa, and an additional role in oocyte maturation or as a maternal effect gene. Progenetic hydrozoans such as *Podocoryna’s* close relative, *Hydractinia*, exhibit a truncation of somatic development and a relatively early maturation of the germline. Evolutionarily, lineages exhibiting progenesis might have undergone relaxed selection on mechanisms controlling somatic development of the medusae, resulting in the loss of *Tlx*.

The phylogenetic distribution and expression pattern suggests that *Tlx* is intimately tied to medusa development. Given that the presence of *Tlx* appears to be ubiquitous in bilaterians, *Tlx* was likely present in the last common ancestor of Bilateria and Cnidaria and was secondarily lost multiple times independently in cnidarians, including at the base of Anthozoa, Endocnidozoa and independently several times in Hydrozoa. The ancestral presence of the *Tlx* gene in Cnidaria, in conjunction with the striking correlation of the gene *Tlx* with the presence of the medusa stage in medusozoans, and its apparent role in medusa development, suggests the possibility of an ancestral metagenetic life cycle in Cnidaria. That is, *Tlx* and medusae could have been present in the ancestor of Cnidaria and lost together multiple times in cnidarian evolution, including at the base of Anthozoa, Endocnidozoa and multiple times in Hydrozoa. An alternative explanation is that *Tlx* could have had an unknown ancestral function in the last common ancestor of Cnidaria and Bilateria and was rapidly exapted in medusozoans for medusa development and/or medusa specific structures or cell types. In this scenario, the ancestral function is no longer necessary in anthozoans and other cnidarians that lack *Tlx*. A third explanation is that *Tlx* has an ancestral function in the development of medusa structures in a cnidarian ancestor that exhibits both polypoid and medusoid features. In that scenario, the emergence of a discrete polyp stage in anthozoans, reduction of the ancestral body plan in parasitic endocnidozoans, and the uncoupling of these features into two discrete generations in medusozoan could have been responsible for the pattern of loss and maintenance of *Tlx* respectively. Further investigations of *Tlx* function in medusozoans and in early diverging bilaterians may help clarify the ancestral role of *Tlx* in Cnidaria.

## Materials and Methods

### Phylogenetic Methods

Amino acid sequence alignments were carried out with Muscle (Edgar, 2004) using default parameters and manually refined on MEGA7 (Kumar et al., 2016). The EH1 domain, N-terminal arm and homeodomain of TLX were aligned with the homeodomain of NK-L members NK6 and HEX, totaling 82 taxa and 83 characters. Phylogenetic analyses were conducted respectively under Mrbayes 3.2.7 (Ronquist and Huelsenbeck, 2003) and RaxML (Stamatakis, 2014), under the model LG+I+G with 4 discrete gamma categories as selected byModeltest-NG (based on BIC), on CIPRES portal (Miller et al. 2011). For the Bayesian phylogenetic analysis, 4 runs and 6 Markov chains were generated, and the analysis was run for 5 million generations with a 25% burn-in. The posterior probabilities, as well as the final topology come from a majority consensus of the sampled trees. For the Maximum likelihood phylogenetic analysis, support values were evaluated by non-parametric bootstraps (1000 replicates). Bootstrap support values were reported when above 50 and posterior probabilities ranging from 0.90 to 1 are indicated at the node in respect to the color coding (Figure 1b).

### Ancestral character state reconstruction of *Tlx* and the gonophore in Medusozoa

The MP ancestral character state reconstruction of the presence of *Tlx* in genomic samples and the gonophore were conducted under Mesquite v3.61 (Maddison and Maddison, 2007) on a pruned tree from Cartwright and Nawrocki, 2010, re-rooted with Acraspeda. Dollo parsimony was imposed for *Tlx* ancestral character reconstruction as convergent evolution or horizontal gene transfer seemed unlikely. The reconstruction of the gonophore ancestral state was conducted under Wagner parsimony; characters were unweighted and unordered. Character states were coded as such: no gonophore, sporosacs, cryptomedusoids, eumedusoids, and medusa. In *Ectopleura larynx*, *Tlx* could not be amplified by degenerate PCR but was found in publicly available transcriptomic data, and thus was coded as present (only one case, *Ectopleura larynx*). Within siphonophores three different structures have been proposed to be homologous to the hydromedusa. The eudoxid is a sexual free-living individual, the nectophore is an asexual swimming zooid exhibiting a velum, radial and circular canals, and the gonophore is the sexual zooid that may or not exhibit medusa-like structures depending on lineages (see Dunn et al., 2005). Here the presence of either nectophores or eudoxids was coded as eumedusoid.

### Cnidarian genomes and transcriptomes assembly

Most of the genome and transcriptome sequences were obtained from the NCBI sequence read archive (SRA) (Table S1) and required assembly in order to screen for *Tlx*. Each library was trimmed of low-quality reads and adapters using fastp (Chen et al., 2018). For those transcriptomes from different libraries, filtered reads were combined into a single dataset followed by *de novo* transcriptome assembly using Trinity v2.8.5 (Grabherr et al., 2011). Genome assemblies were carried out using Spades v3.13.1 (Bankevich et al., 2012). Genomes that did not require assembly were obtained from NCBI or from unpublished work shared by collaborators. The source of all the genomes and transcriptomes used in this study can be found in Table S1.

### *In silico* search of *Tlx* and phylogenetic analyses

Potential orthologs of *Tlx* were identified through reciprocal blasts of TLX amino acid sequences (tblastn, e-value cut-off set to 10^-80^ and 10^-10^) from *Chironex fleckeri, Calvadosia cruxmelitensis, Aurelia aurita, Clytia hemisphaerica, Podocoryna carnea, Agalma elegans*, and *Craspedacusta sowerbii* against cnidarian transcriptomes and genomes. Phylogenetic methods are outlined in the Supplementary Information.

### Degenerate PCR screening of *Tlx* from cnidarian genomic DNA

Degenerate primers in the second exon of *Tlx* (Forward 5’-GGNCAYCCNTAYCAVAGC/MGNGC-3’, Reverse 5’-GTKCKHCKRTTYTGAWACCA-3’) were designed from 8 medusozoan species: *Chironex fleckeri, Calvadosia cruxmelitensis Aurelia aurita, Clytia hemisphaerica, Podocoryna carnea, Agalma elegans* and *Craspedacusta sowerbii*. The primers span the N-terminal arm to the C-terminal end of TLX homeodomain, for a total expected amplicon length of 196 bp. Degenerate PCR were carried out using OneTaq 2X Master Mix Standard Buffer, according to manufacturer instructions and with an annealing temperature of 40° C. Amplification products were electrophoresed in a 1.2% agarose gel to assess the presence of the amplicon. The amplifiability of the genomic samples was assessed using 16S degenerate primers. Ten amplicons were selected at random, cloned into the pCR4-TOPO plasmid and sequenced to confirm TLX identity.

### Bayesian correlation analysis of the presence of *Tlx* and the medusa stage

Character coding and ancestral state reconstruction methods are outlined in the Supplementary Inforrmation. Correlation analyses between the presence of *Tlx* and medusa stage was performed using BayesTraits v3 (Pagel et al., 2004), imposing irreversibility for *Tlx* by setting transition rate for regains of *Tlx* to zero (q_13_=0, q_24_=0). The presence of *Tlx* and the medusa stage were coded as binary characters. The presence of sporosacs and cryptomedusoids as well as no gonophores were coded as absent and the eumedusoids and fully developed medusae were coded as present. Although eumedusoids do not feed, they have nearly all other medusa components and thus were treated as medusae. The statistical support for the correlation analysis was carried out by computing the marginal likelihood of the two alternative models, independence and dependence of *Tlx* with the medusa stage. The calculation of the Log Bayes Factor was performed and interpreted as recommended by the BayesTraits manual. According to the Bayestraits manual logBF can be interpreted as such: logBF <2 Weak evidence, logBF >2 Positive evidence, 5 ≤ logBF ≤ 10 Strong evidence, logBF >10 Very strong evidence.

### Animal care

*P. carnea* colonies were grown on microscope slides contained in slide racks and kept in artificial seawater (REEF CRYSTALS, Aquarium Systems) in a 7L Kreisel tank at room temperature (~18°C) with a salinity of 29 ppt. Male and female colonies were kept in separate tanks. Colonies were fed two-day old *Artemia* nauplii twice a week and blended mussels once a week. Unfed one and three day old released medusae were collected. Prior to every experiment, *P.carnea* colonies were starved for four days. Animals were relaxed for 30min by addition of menthol crystals (1mg/ml) to the medium and fixed after two medium changes.

### Probe synthesis and *In situ* hybridization of *Tlx* in *P.carnea*

The sequence for *Tlx* transcript was recovered from a newly assembled transcriptome of *P. carnea. Tlx* was amplified from medusae cDNA using the following PCR primers: *P. carnea* forward 5’-GAAAGATAAACACGAAAAAGAAACGG-3’ and reverse 5’-TCCGGAACTTCATTACTCGCTGTTGC-3’ for an expected amplicon length of 528 bp. Amplicons were cloned using the Invitrogen pCR4-TOPO-TA Cloning Kit and sequenced using M13 forward and reverse primers. Sense and antisense DIG labeled riboprobes were synthesized from clones using the Invitrogen T7/T3 Megascript kit. *In situ* hybridization (ISH) protocol was adapted from Gajewsky et al, 1996. Animals were fixed in ice cold fix (3.7%PFA and 0.25% glutaraldehyde in 1X PBS). Hybridization was carried out at 50°C for 18 hours with a probe concentration of 1 ng/μl. DIG labeled riboprobes localization was detected by immunostaining with anti-DIG-Fab-AP (ROCHE) and NBT/BCIP.

### *De novo* transcriptome assembly for *P. carnea*

For the *Podocoryna carnea de novo* transcriptome assembly used in this study, planula and early budding polyp libraries were sequenced and the sequences submitted to NCBI Sequence Read Archive (BioProject ID PRJNA744579) in addition to updated libraries for non-reproductive polyps, budding polyps and medusae (BioProject ID PRJNA245897), and used along with previously generated *P. carnea* libraries described in Sanders and Cartwright, 2015. Stage 2 planulae (elongated, swimming, non-competent) and stage 1-2 early budding polyps, see Frey (1968), were flash frozen and sent to KUMC-GSF for RNA library preparation using the TruSeq RNA Sample Preparation Kit (BoxA). All libraries were 100 bp paired-end with an average insert size of 170 bp. Libraries were then barcoded, pooled, and multiplexed on a single lane of an Illumina NovaSeq 6000 S1 flow cell at KUMC-GSF. Low-quality reads were trimmed and adapters using fastp (Chen et al., 2018). Reads from all libraries but the planula ones were mapped to the draft genomes of their respective strains of *Podocoryna carnea* (Chang and Baxevanis, pers. comm.) using STAR (Dobin et al, 2013). Uniquely mapped reads (61.28%) and reads mapping on multiple loci (20.05%) were kept for *de novo* assembly. The *de novo* transcriptome assembly was produced with Trinity v2.8.5 (Grabherr et al., 2011), yielding 472366 transcripts total with an average length of 714.62 and a G+C content of 36.76%. Transcripts were blasted against a *Mus musculus* transcriptome dataset (GCA_000001635.9 GRCm39) with an e-value threshold of 1.e^-100^, and best hits were removed from the assembly. The longest ORFs from the transcriptomes were predicted using Transdecoder version 5.5.0 (http://transdecoder.github.io), duplicate sequences and isoforms were removed by clustering sequences with a 95% identity threshold using CD-HIT version 4.8.1 (Fu et al, 2012) and a BUSCO analysis (Simão et al, 2015) against the metazoan database (metazoa_odb10) was performed to assess the completeness of the transcriptome, estimating a 96.2% completeness of the transcriptome (82.9% single copy Buscos and 13.3% duplicated Buscos), 1.5% of fragmented Buscos and 2.3% of missing Buscos on a total of 954 BUSCO markers.

### Differential expression analysis in the life cycle of *P. carnea, C. hemisphaerica and A. coerulea* and *Tlx* qPCR validation in *P.carnea*

The differential expression analyses were carried out on the transcriptome of *Clytia hemisphaerica* (http://marimba.obs-vlfr.fr), *Aurelia coerulea* (https://davidadlergold.faculty.ucdavis.edu) and the *de novo* transcriptome assembly of *Podocoryna carnea*. The reads quantification was performed at the gene level, quantification combining isoforms, using RSEM (Li and Dewey, 2011). Estimated counts, Fragments per Kilobase Million (FPKM) and Transcripts per Million (TPM) values were generated and differential expression of *Tlx* for the three species was analyzed on these three metrics using Ebseq (Leng et al. 2013). The differential expression analysis was performed on the main developmental stages of the life cycle for the three species, planulae (binning the three planulae stages for *Clytia*), non-reproductive polyp (respectively, gastrozooid, scyphostoma and non-reproductive polyp), reproductive polyp (respectively, gonozooid, early and late budding polyps, early and late strobila) and medusa (respectively, ephyra, juvenile and mature). The relative expression of *Tlx* was validated in *Podocoryna carnea* through RT-qPCR. Tissues from planulae (7 biological replicates) non-reproductive polyps (8 biological replicates), budding polyps (8 biological replicates) and released medusae (8 biological replicates) were homogenized in Trizol and incubated for 15 min. Samples were then combined with 0.5 volume of chloroform, mixed, incubated for 3 min and then spun down at 4C for 15min. The supernatant was mixed with one volume of 70% ethanol and RNA extraction was carried out using a Qiagen RNAeasy Micro Kit. cDNA synthesis was carried out using Superscript IV and a primer mixture of random hexamers and oligo-dT. cDNA was quantified with Qubit and the qPCR was performed using PowerUP SYBR Green, using the following primers (*Ef1* forward 5’-TTGCCACCTCAACGACCATC-3’, *Ef1* reverse 5’-TACCGACTGGCACTGTTCCA-3’ and *Tlx* forward 5’-CAGAGCCCCACCGAAAAGAA-3’, *Tlx* reverse 5’-ATTCCTTGGCCACACGCAAT-3’).

## Acknowledgments

We thank A. Baxevanis and E.S Chang for providing draft genome sequences, A. Klompen for help in data collection and R. Helm and Catriona Munro for providing access to transcriptome sequences. Genome sequencing services was provided by the University of Kansas Medical School Genome Sequencing Center. Computing support was provided by the University of Kansas Center for Research Computing. This work was supported by the National Science Foundation (NSF) Division of Environmental Biology 095357 the University of Kansas General Research Fund, and the University of Kansas Research Excellence Initiative (to PC) and support to PC while serving at the National Science Foundation MT was supported by the Chancellor’s Doctoral Fellowship at the University of Kansas and received research support from the KU Graduate Research Fund.

## Notes

**Competing Interest Statement:** The authors declare no conflict of interest.

### Competing Interest Statement

The authors have declared no competing interest.

## References

1. Allman, G. J. (1871). A Monograph of the Gymnoblastic Or Tubularian Hydroids. Ray Society.

2. Bankevich, A et al. (2012). SPAdes: A New Genome Assembly Algorithm and Its Applications to Single-Cell Sequencing. Journal of Computational Biology, 19(5), 455– 477.

3. Boero, F., Bouillon, J., & Piraino, S. (1992). On the origins and evolution of hydromedusan life cycles (Cnidaria, Hydrozoa). In R. Dallai (Ed.) Sex Origin and Evolution., Selected Symposia and Monograph UZI 6. Mucchi, 59–68.

4. Boero, F., Bouillon, J., Cicogna, F., & Cornelius, P. (1987). Inconsistent evolution and paedomorphosis among the hydroids and medusae of the Athecatae/Anthomedusae and the Thecatae/Leptomedusae (Cnidaria, Hydrozoa). Modern trends in the systematics, ecology, and evolution of hydroids and hydromedusae, 229–250.

5. Bouillon, J., Gravili, C., Gili, J. M., & Boero, F. (2006). An Introduction to Hydrozoa. Mémoires Du Muséum National d’Histoire Naturelle, 194, 1–591.

6. Cartwright, P., & Nawrocki, A. M. (2010). Character Evolution in Hydrozoa (phylum Cnidaria). Integrative and Comparative Biology, 50(3), 456–472.

7. Chang, E.S et al. (2015) Genomic insights into the evolutionary origin of Myxozoa within Cnidaria. Proc Natl Acad Sci USA 112(48):14912–14917.

8. Chen, S., Zhou, Y., Chen, Y., & Gu, J. (2018). fastp: An ultra-fast all-in-one FASTQ preprocessor. Bioinformatics, 34(17), i884–i890.

9. Cunningham, C. W., & Buss, L. W. (1993). Molecular evidence for multiple episodes of paedomorphosis in the family Hydractiniidae. Biochemical Systematics and Ecology, 21(1), 57–69.

10. Derelle, R., & Manuel, M. (2007). Ancient connection between NKL genes and the mesoderm? Insights from Tlx expression in a ctenophore. Development Genes and Evolution, 217(4), 253–261.

11. Dobin, A. et al. (2013). STAR: Ultrafast universal RNA-seq aligner. Bioinformatics, 29(1), 15–21.

12. Dunn, C. W. (n.d.). The colony-level evolution and development of the siphonophora (Cnidaria, Hydrozoa) [Ph.D., Yale University].

13. Edgar, R. C. (2004). MUSCLE: Multiple sequence alignment with high accuracy and high throughput. Nucleic Acids Research, 32(5), 1792–1797.

14. Frank, U., Nicotra, M. L., & Schnitzler, C. E. (2020). The colonial cnidarian Hydractinia. EvoDevo, 11(1), 7.

15. Frey, J. (1968). Die Entwicklungsleistungen der Medusenknospen und Medusen vonPodocoryne carnea M. Sars nach Isolation und Dissoziation. Wilhelm Roux’ Archiv fur Entwicklungsmechanik der Organismen, 160(4), 428–464.

16. Fu, L et al. (2012). CD-HIT: Accelerated for clustering the next-generation sequencing data. Bioinformatics, 28(23), 3150–3152.

17. Gajewski, M., Leitz, T., Schloßherr, J., & Plickert, G. (1996). LWamides from Cnidaria constitute a novel family of neuropeptides with morphogenetic activity. Roux’s Archives of Developmental Biology, 205(5), 232–242.

18. Galliot, B. (2012). *Hydra*, a fruitful model system for 270 years. International Journal of Developmental Biology, 56(6-7–8), 411–423.

19. Goette, A. (1916). Die Gattungen *Podocoryne, Stylactis* und Hydractinia. Zoologische Jahrbücher, 39, 443–510.

20. Gould, S. J. (1985). Ontogeny and Phylogeny. Harvard University Press.

21. Grabherr, M. G et al. (2011). Trinity: Reconstructing a full-length transcriptome without a genome from RNA-Seq data. Nature Biotechnology, 29(7), 644–652.

22. Hincks, T. (1868). A History of the British Hydroid Zoophytes: 1: Text. John Van Voorst.

23. Kamm, K., & Schierwater, B. (2006). Ancient complexity of the non-Hox ANTP gene complement in the anthozoan *Nematostella vectensis*. Implications for the evolution of the ANTP superclass. Journal of Experimental Zoology Part B: Molecular and Developmental Evolution, 306B(6), 589–596.

24. Khalturin, K et al. (2019). Medusozoan genomes inform the evolution of the jellyfish body plan. Nature Ecology & Evolution, 3(5), 811–822.

25. Kojima, T., Tsuji, T., & Saigo, K. (2005). A concerted action of a paired-type homeobox gene, aristaless, and a homolog of Hox11/tlx homeobox gene, clawless, is essential for the distal tip development of the Drosophila leg. Developmental Biology, 279(2), 434–445.

26. Kraus, J. E et al. (2015). Adoption of conserved developmental genes in development and origin of the medusa body plan. EvoDevo, 6(1), 23.

27. Kubota, S. (2000). Parallel, paedomorphic evolutionary processes of the bivalve-inhabiting hydrozoans (Leptomedusae, Eirenidae) deduced from the morphology, life cycle and biogeography, with special reference to taxonomic treatment of Eugymnanthea. Scientia Marina, 64(S1), 241–247.

28. Kumar, S., Stecher, G., & Tamura, K. (2016). MEGA7: Molecular Evolutionary Genetics Analysis Version 7.0 for Bigger Datasets. Molecular Biology and Evolution, 33(7), 1870–1874.

29. Leclère, L et al. (2019). The genome of the jellyfish *Clytia hemisphaerica* and the evolution of the cnidarian life-cycle. Nature Ecology & Evolution, 3(5), 801–810.

30. Leng, N et al. (2013). EBSeq: An empirical Bayes hierarchical model for inference in RNA-seq experiments. Bioinformatics, 29(8), 1035–1043.

31. Lenti, E et al. (2016). Transcription factor TLX1 controls retinoic acid signaling to ensure spleen development. The Journal of Clinical Investigation, 126(7), 2452–2464.

32. Li, B., & Dewey, C. N. (2011). RSEM: Accurate transcript quantification from RNA-Seq data with or without a reference genome. BMC Bioinformatics, 12(1), 323.

33. Maddison, W., & Maddison, D. (2007). Mesquite 2. A Modular System for Evolutionary Analysis. Available online at: http://www.mesquiteproject.org.

34. Masuda-Nakagawa, L. M., Gröer, H., Aerne, B. L., & Schmid, V. (2000). The HOX-like gene Cnox2-Pc is expressed at the anterior region in all life cycle stages of the jellyfish *Podocoryne carnea*. Development Genes and Evolution, 210(3), 151–156.

35. Miller, M. A., Pfeiffer, W., & Schwartz, T. (2011, July). The CIPRES science gateway: a community resource for phylogenetic analyses. In Proceedings of the 2011 TeraGrid Conference: extreme digital discovery (pp. 1–8).

36. Müller, P., Yanze, N., Schmid, V., & Spring, J. (1999). The Homeobox Gene Otx of the Jellyfish *Podocoryne carnea*: Role of a Head Gene in Striated Muscle and Evolution. Developmental Biology, 216(2), 582–594.

37. Murrell, B et al. (2013). FUBAR: A Fast, Unconstrained Bayesian AppRoximation for Inferring Selection. Molecular Biology and Evolution, 30(5), 1196–1205.

38. Namikawa H. (1991). A New Species of the Genus *Stylactaria* (Cnidaria,Hydrozoa) from Hokkaido, Japan. Zoological Science, 8(4), p805–812.

39. Pagel, M., Meade, A., & Barker, D. (2004). Bayesian estimation of ancestral character states on phylogenies. Systematic biology, 53(5), 673–684.

40. Pang, K., & Martindale, M. Q. (2008). Developmental expression of homeobox genes in the ctenophore Mnemiopsis leidyi. Development Genes and Evolution, 218(6), 307–319.

41. Ronquist, F., & Huelsenbeck, J. P. (2003). MrBayes 3: Bayesian phylogenetic inference under mixed models. Bioinformatics, 19(12), 1572–1574.

42. Ryan, J. F et al. (2006). The cnidarian-bilaterian ancestor possessed at least 56 homeoboxes: Evidence from the starlet sea anemone, *Nematostella vectensis*. Genome Biology, 7(7), R64.

43. Sanders, S. M., & Cartwright, P. (2015a). Patterns of Wnt signaling in the life cycle of *Podocoryna carnea* and its implications for medusae evolution in Hydrozoa (Cnidaria). Evolution & Development, 17(6), 325–336.

44. Sanders, S. M., & Cartwright, P. (2015b). Interspecific Differential Expression Analysis of RNA-Seq Data Yields Insight into Life Cycle Variation in Hydractiniid Hydrozoans. Genome Biology and Evolution, 7(8), 2417–2431.

45. Simão, F et al. (2015). BUSCO: Assessing genome assembly and annotation completeness with single-copy orthologs. Bioinformatics, 31(19), 3210–3212.

46. Stamatakis, A. (2014). RAxML version 8: A tool for phylogenetic analysis and post-analysis of large phylogenies. Bioinformatics, 30(9), 1312–1313.

